# Polyamine depletion inhibits norovirus infection by blocking virus-induced apoptosis

**DOI:** 10.1101/2025.09.14.676151

**Authors:** Maryna Chaika, Heike Laschin, Marina Pekelis, Carmen Mirabelli, Sandra Niendorf, Christiane E. Wobus, Stefan Taube

## Abstract

Noroviruses are non-enveloped, positive-sense RNA viruses in the family *Caliciviridae*. Human noroviruses (HNoVs) are the leading cause of viral gastrointestinal disease. Despite their clinical relevance, therapeutic strategies for norovirus infections remain limited, partly due to challenges in cultivating HNoV and limited understanding of virus-host interactions. Polyamines (PAs) are small polycationic metabolites derived from amino acid metabolisms and are essential for diverse cellular functions and important host factors for many RNA viruses. However, their role in norovirus infection has remained elusive. Here, we demonstrate that PAs are critical for productive infection for both HNoV and murine norovirus (MNV), underscoring a conserved and functionally significant dependency across species. Functional analysis in MNV-infected murine macrophages showed that PA depletion did not significantly impair virus attachment and viral genome replication. However, it resulted in a marked reduction in infectious virus titers following the first replication cycle, coinciding with a loss of virus-induced apoptosis and impaired virion release. Exogenous supplementation with two metabolites of the PAs biosynthetic pathway (spermine or spermidine), or induction of apoptosis using a PI3-kinase inhibitor, restored both viral titers and apoptosis in PA-depleted macrophages. In human intestinal enteroids (HIEs), depletion of PAs completely ablated HNoV infection, which was partially rescued adding exogenous PAs. Collectively, our findings establish PAs as a host dependency factor that facilitates norovirus infection, promoting virus-induced apoptosis and egress in the murine system. Furthermore, these results point to PAs metabolism as a potential therapeutic target for antiviral intervention against HNoV infections.

**Importance:** HNoV are a major cause of acute viral gastroenteritis worldwide, yet the development of effective antivirals and vaccines has been hampered by the lack of small-animal models and robust *in vitro* systems to propagate the virus. MNV shares key features with HNoV and serves as a valuable surrogate for studying norovirus biology. In this study, we identify host-derived PAs as essential and conserved metabolites required for productive infection for both HNoV and MNV. PAs exert multifactorial effects on norovirus infection, particularly by facilitating virus-induced apoptosis and promoting virion release in MNV-infected cells. Notably, PA depletion using difluoromethylornithine (DFMO), a licensed and well-tolerated inhibitor of PA synthesis, effectively suppressed norovirus infection. Our findings highlight the potential of repurposing DFMO as a host-targeting antiviral, especially for the treatment of chronic norovirus infections in immunocompromised individuals. This work provides new insight into norovirus-host interactions and proposes a readily translatable therapeutic avenue.

## Introduction

Noroviruses are non-enveloped, positive-sense single-stranded RNA (+ssRNA) viruses belonging to the *Caliciviridae* family (1). Human norovirus (HNoV) is a leading cause of viral gastroenteritis worldwide, characterized by symptoms ranging from acute vomiting to diarrhea (2). Although HNoV infection is typically self-limiting in healthy individuals, it can cause prolonged illness and chronic viral shedding in immunocompromised patients, with the potential for severe or even fatal outcomes (3). Despite its global health impact, no licensed antivirals or prophylactic vaccines are currently available for HNoV (4). Progress in therapeutic development has been hindered by the historical lack of suitable models to support robust HNoV replication *in vitro* (*5*). While histo-blood group antigens (HBGAs) have long been recognized as key host factors, they are not sufficient for productive infection (6). A major breakthrough in the field was the establishment of stem cell-derived human intestinal enteroid (HIE) cultures, which support HNoV replication from human stool isolates (7, 8). This system has enabled critical advances in our understanding of norovirus pathogenesis and has facilitated the first antiviral drug screens (9).

Murine norovirus (MNV), discovered in 2003, remains the only norovirus that can be productively cultivated in cell culture (10). MNV shares numerous molecular, structural, and biological characteristics with HNoV, making it a valuable surrogate model for studying HNoV biology (11). A key distinction, however, is MNV’s use of the murine CD300lf/ld protein as a *bona fide* entry receptor (12, 13), which currently has no known functional homolog in human HNoV infection (6).

Like other enteric viruses, MNV utilizes both lytic and non-lytic egress mechanisms, influencing viral pathogenesis and dissemination (14). Acute strains, such as MNV-1, induce lytic infection in myeloid-derived cells, including RAW264.7 macrophages, BV-2 microglial cells (15), as well as primary monocyte-derived cells (16). In these models, infection causes pronounced cytopathic effects (CPE). In immunocompetent mice, MNV-1 infection is self-limiting and typically resolves within seven days post-infection (17). In contrast, persistent strains such as CR3 or CR6 establish long-term infection by targeting specialized intestinal epithelial cells known as tuft cells, where they evade immune clearance through suppression of host antiviral responses (18–20).

Apoptosis, a non-immunogenic form of programmed cell death (PCD), is a common host defense mechanism against infection, characterized by the orderly disassembly of cells into membrane-bound fragments known as apoptotic bodies (21). Hallmark features of apoptosis include activation of caspases, cleavage of DNA repair enzymes such as Poly (ADP-ribose) polymerase (PARP) by host caspases, resulting in a loss of the cellular DNA damage response (22), as well as downregulation of pro-survival proteins like survivin (23) and Mcl-1, and the externalization of phosphatidylserine (PS) to the outer leaflet of the plasma membrane (24). The phosphoinositide 3-kinase (PI3K)-Akt-mTOR pathway is frequently hijacked by RNA viruses to modulate apoptosis and enhance viral replication (25).

While many viruses have evolved mechanisms to inhibit apoptosis and prolong host cell survival (26), noroviruses strategically modulate the intrinsic apoptotic pathway to facilitate their replication and dissemination (27, 28). MNV NS3 carries a mitochondrial localization signal and an MLKL-like membrane-disruption domain that drives cell death through mitochondrial targeting (28). During early stages of infection, MNV suppresses apoptosis i.e. via activation of the PI3K/Akt signaling pathway (14). This early inhibition of apoptosis allows sufficient time for viral replication and assembly and influences the balance between intracellular retention and release of virions. At later stages, MNV egress becomes associated with virus-induced apoptosis, mediated by host proteases, such as caspases and cathepsins (29), dysregulation of pro-survival BCL-2 family (30), and expression of an immunomodulatory protein VF1 (31, 32). Additionally, caspase-mediated cleavage of NS1/2 further enhances intrinsic apoptosis and is essential for the intestinal epithelial cell tropism of persistent MNV strains (33).

In contrast, the mechanisms by which HNoV regulates apoptosis are less well understood. However, as for MNV, the HNoV non-structural protein NS3 (NTPase) has been shown to induce apoptosis, as evidenced by caspase activation and PARP cleavage (27, 34). Notably, other viral proteins such as NS1/2 (N-terminal domain) and NS4 (P22) physically interact with NS3 and enhance its pro-apoptotic activity (35), suggesting that coordinated modulation of host cell death pathways may be a conserved feature of norovirus infection.

Viruses rely on host factors, to complete their infection cycle; among these, metabolites, such as polyamines (PA)s have emerged as essential regulators of viral replication and pathogenesis (36, 37). PAs play a crucial role in maintaining intestinal barrier integrity by regulating epithelial cell proliferation, migration, and tight junction stability (38). PAs are also crucial for cellular processes, such as cell-cycling, nucleic acid binding, apoptosis regulation and membrane fluidity (39–43). PA are also involved in translation regulation, through hypusination, where PAs are covalently attached to the eukaryotic translation initiation factor eIF5A (44). Hypusination then promotes expression of cholesterol synthesis genes by supporting SREBP2 synthesis.

Mammalian cells exclusively synthesize biogenic PAs: e.g., putrescine, spermidine, and spermine, from an ornithine precursor through ornithine decarboxylase (ODC1) (36) (**Figure 1a**). PA depletion is associated with decreased expression of caspases and pro-apoptotic Bax/Bad proteins (45) and leads to cell survival upon stimulation of intestinal epithelial cells with extrinsic apoptotic signal (46).

**Figure 1.**
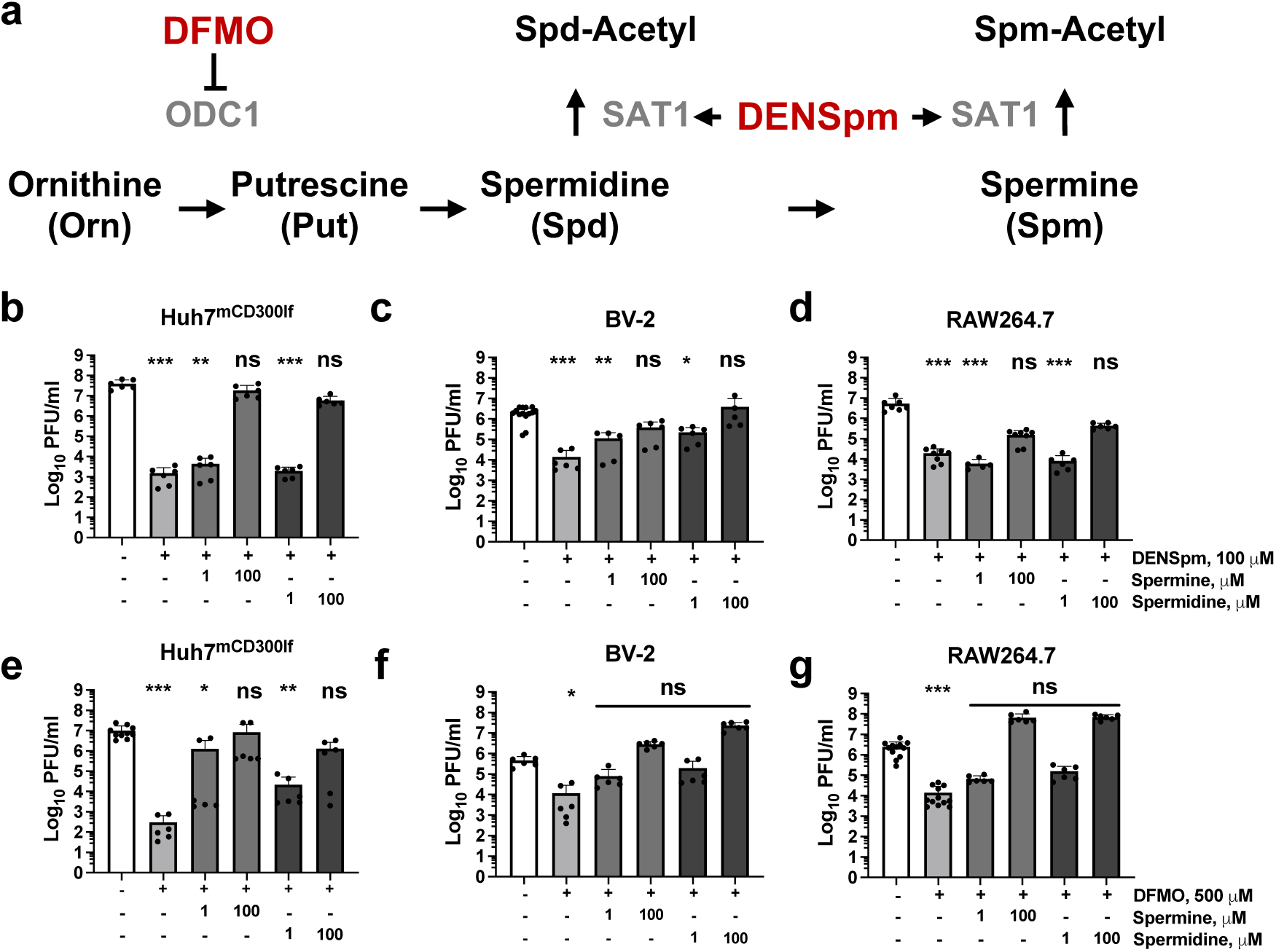
PA depletion restricts MNV infection in immune and epithelial cells. Polyamines Biosynthesis and Catabolic Pathway **(a)**. Huh7^mCD300lf^ **(b, e)**, BV-2 **(c, f)** and RAW264.7 **(d, g)** were pretreated with 100 μM DENSpm for 24 h **(b-d)** or 500 μM DFMO for 96 h **(e-g)**. Cells were inoculated with MNV-1 at MOI 0.1 and for 1 h at 4°C. The inoculum was removed and replaced with culture medium (-) or culture medium supplemented with DENSpm, DFMO with or without 1 or 100 μM of spermine or spermidine. Samples were lysed 24 hours post infection (hpi) and virus titers were determined by plaque assay. Data represent means ± standard deviation (SD) of three independent biological assays with two technical repeats. Individual data points are shown. Statistical significance was assessed using the Kruskal–Wallis test compared to non-treated/infected: ns, not significant; *p≤0.05; **p≤0.01; ***p≤0.001.

PAs are known as host factors for a broad range of RNA and DNA viruses (36, 37), by playing a role at different steps of viral infection. For example, Coxsackievirus B3 (CVB3), a picornavirus, requires PA for cell entry (44), while Zika virus (ZV) and other Flaviviruses require PAs for genome replication (47, 48). Bunyaviruses, like Rift Valley Fever Virus (RVFV), incorporate PAs into virions and rely on PAs for production of infectious particles (49, 50).

Here, we show that depletion or degradation of PAs inhibits infection of both HNoV and MNV, highlighting a conserved requirement for PAs across norovirus genogroups. Notably, PA depletion protected MNV-infected macrophages from virus-induced cell death for up to two days post-infection. PA depletion specifically blocked MNV-induced PARP cleavage, a hallmark of apoptosis and was restored by the PI3-kinase inhibitor LY294002 (LY). Our findings corroborate the importance of the crosstalk between norovirus infection and apoptotic pathways and show a role of PAs in norovirus-induced apoptosis.

## Materials and Methods

### Cell culture

Murine microglial cell line BV-2, murine macrophage cell line RAW264.7, and recombinant human hepatoma cell line Huh7, stably expressing murine CD300lf (Huh7^mCD300lf^) (51) were used. BV-2 cells were maintained in DMEM (Dulbeco’s Modified Eagle Medium) supplemented with 5 % fetal calf serum (FCS), 1 % penicillin/streptomycin (P/S) (Capricorn Scientific), 1 % L-glutamine (L-Glut) (C-C-Pro) and 1 % non-essential amino acids (NEAA) (Capricorn Scientific). RAW264.7 and Huh7^mCD300lf^ were maintained in DMEM supplemented with 10 % FCS, 1 % P/S, 1 % L-Glut and 1 % NEAA. For passaging, RAW264.7 and BV-2 cells were detached by scraping and Huh7^mCD300lf^ by trypsinization using 0.05% Trypsin-EDTA (Thermo Fisher Scientific). All cells were incubated at 37°C, 5 % CO_2_, 95 % relative humidity (RH).

### Preparation of MNV virus stocks

The MNV-1 (GV/MNV1/2002/USA) CW1 isolate was generated by reverse genetics with minimal passaging (51) and used up to passage 4. Wild type CR3 (GV/CR3/2005/USA) was used up to passage 4, since plaque isolation. Virus stocks were prepared by infecting BV-2 cells at a multiplicity of infection (MOI) ∼ 0.1 until strong cytopathic effect was visible, usually ∼ 24 hours post-infection (hpi). Flasks were freeze-thawed two times and lysate centrifuged at 4,500×g for 10 min at 4°C to remove cell debris. The clarified supernatant was precipitated by 40 % ammonium sulfate, overnight, rotating at 4°C. Precipitated virus was pelleted by centrifugation at 10,000×g for 15 min at 4°C. Virus was resuspended in 100 mM sodium phosphate buffer pH 7.4, sterile-filtered (0.22 µm pore size). Titered aliquots were stored at -80°C.

### Chemical inhibitors

Difluoromethylornithine hydrochloride hydrate (DFMO, also known as Eflornithine) (Tebu-Bio, #T2593), and N1, N11-Diethylnorspermine tetrahydrochloride (DENSpm) (Bio-Techne, #0468) were prepared in ddH_2_O as 50 mM and 2.5 mM stocks, respectively, and stored in aliquots at -20°C. LY294002 (LY) hydrochloride (Tebu-Bio, #T4079) was prepared in dimethyl sulfoxide (DMSO) (Carl Roth, #A994.2) as 5 mM stocks, and stored in aliquots at -20°C. Staurosporine (BioMol, #LKT-S7600) was prepared in DMSO as 1 mM stocks, and stored in aliquots at -20°C. Spermine (Cayman Chemicals, #18041) and spermidine (Cayman Chemicals #14918) were prepared as 1 mM stocks in DMEM supplemented with 1 % P/S and 1 % L-Glut, and stored short term at 4°C.

### PA depletion/degradation and MNV infection

For PA depletion, cells were treated with 500 µM DFMO at ∼60 – 70 % confluency for 4 days. At day three, cells were re-seeded in 24-well tissue culture plates at 4×10^5^ cells/well for RAW264.7 and at 2×10^5^ cells/well for BV-2 and Huh7^mCD300lf^ cells and freshly supplemented with DFMO containing medium for the final day of treatment. For PA degradation, cells were treated with DENSpm at 100 µM after plating and incubated overnight for an 18 h treatment. For MNV infections, cells were inoculated with MNV diluted in cold PBS at indicated MOI and incubated for 1 h on ice. Virus inoculum was removed, and cells were washed three times with ice-cold PBS after which warm culture medium plus the indicated supplements and incubated at 37°C in 5 % CO_2_ at 95 % RT as indicated. After incubation, cells and/or supernatant were lysed by freeze-thawing twice and titered as indicated, either by plaque assay or endpoint dilution. For protein analysis by sodium dodecyl sulfate (SDS)-polyacrylamide gel electrophoresis (PAGE) and western blotting, RAW264.7 cells were seeded in 6-well cell culture plates at 1×10^6^ cell/well before infection the following day as described above. At the experimental endpoint, the supernatant was removed, and cells were collected in an ice-cold lysis buffer (PBS, 100 mM NaCl, 1 % Triton-X, pH 8.0) before protein extraction.

### Plaque assay

Infectious virus titers were measured using plaque assay as previous described (52). Briefly, BV-2, RAW264.7, or Huh7^mCD300lf^ cells (51) were plated in 6-well tissue culture plates one day prior to infection at 5×10^5^, 5×10^6^ or 2×10^5^ cells/well, respectively. As BV-2 cells were less adherent to our cell culture plastic, end point dilutions were the preferred method for this titration. Infectious samples were serially diluted 1:10 in a total volume of 1.25 ml (1,125 µL PBS or sodium phosphate buffer, 125 µL sample). Cell culture medium was aspirated, and cells were infected in duplicate with 500 µL of the respective diluted virus per well. Inoculated plates were incubated at 37°C, 5 % CO_2_, 95 % RH for 1 h. Virus was aspirated and replaced with an overlay medium (1.2 % Avicel^®^ (IMCD Deutschland GmbH, #RC-581NF), 1x MEM (Thermo Fisher Scientific, #61100087), 5 % FCS; 1 % P/S, 1 % L-Glut). Plates were incubated at 37°C, 5 % CO_2_, 95 % RH for 48 hrs. The overlay medium was aspirated, cells were washed with PBS and stained with 1 mg/ml Erythrosine B (Sigma-Adrich, #198269) in PBS.

### Endpoint titration determining tissue culture infectious dose 50 (TCID_50_)

Infectious virus titers were measured by endpoint titration (53), as an alternative to the preferred plaque assay. BV-2 cell were the preferred cell line, as only 1×10⁴ cells/96-well were needed and no trypsinization was required. Cells were seeded in flat-bottom 96-well tissue culture plates and used immediately after cell adherence. Virus containing samples were serially diluted using 180 μL media and of 20 μL virus sample. Plates were incubated for 3 to 4 days at 37 °C, 5 % CO₂, and 95 % RH. After incubation, wells were scored for CPE by light microscopy. Viral titers were calculated as TCID₅₀/ml using the Reed–Muench method (54, 55).

### qRT-PCR based MNV-1 attachment assay

Mock- or DFMO-treated Huh7^mCD300lf^, BV-2, or RAW264.7 cells were seeded one day prior to infection in 6-well tissue culture plates (2×10⁵, 5×10⁵, 5×10^6^ cells/well, respectively). The following day, virus inoculum was prepared to an MOI 5 by diluting MNV-1.CW1 stock in PBS (pH 7.4) supplemented with spermine and/or DFMO. A total of 500 µl inoculum was added to each well and incubated on ice for 1 h to allow virus attachment. Following incubation, the inoculum was aspirated, and cells were washed three times with ice-cold PBS (pH 7.4) to remove unbound virus. After the final wash, wells were aspirated completely, and cell monolayers were lysed in 500 µL TRIzol™ reagent (Zymo Research Europe GmbH). Genome equivalents (GEs) were determined as previously described (51) and normalized to the total amount of RNA extracted. Briefly, total RNA was extracted using the Direct-zol RNA MiniPrep kit (Zymo Research Europe GmbH) according to the manufacturer’s instructions. RNA concentration was determined using a NanoDrop 2000 spectrophotometer (Thermo Fisher Scientific), assessing A260/280 and A260/230 ratios. GEs were determined using the Luna Universal Probe One-Step RT-qPCR Kit (NEB) with primers and a probe targeting the NS1/2 in ORF1 (Forward: 5′-GTGCGCAACACAGAGAAACG; Reverse: 5′-CGGGCTGAGCTTCCTGC; Probe: 5′-[6-FAM]-CTAGTGTCTCCTTTGGAGCACCTA-[BHQ1]). A synthetic dsDNA fragment (gBlock™, Integrated DNA Technologies IDT) corresponding to the amplified cDNA was used to generate a standard curve for quantification of genome eq.

### Antibodies

Antibodies against PARP and Caspase-3 (Cell Signaling Technologies, #9542T, #9662S respectively) were diluted 1:1,000 for protein detection during western blotting. Beta–actin (Bio-Teche, #MAB8929-SP) was used at a final concentration of 0.02 µg/ml. The broadly specific norovirus VP1-specific monoclonal antibody TV19 (56), was used at 1:2,000 dilution (kindly provided by Peter Sander, R-Biopharm AG, Germany). Secondary antibodies goat anti-rabbit-HRP IgG (Dianova, #111-045-003) and goat anti-mouse (Dianova, #111-035-003) were diluted 1:3,000 during western blotting.

### Protein extraction

Cells were scraped in lysis buffer (PBS, 100 mM NaCl 1 % Triton-X, pH 8.0) from wells using a sterile cell scraper and transferred into sterile centrifuge tubes. To ensure maximum cell lysis, tubes were incubated on ice for 10 to 15 min. Lysed cells were centrifuged at 4°C, 16,000×g for 20 min. The cleared lysate was transferred into fresh tubes and mixed (1:2) with sample loading buffer (62.5 mM Tris, 10 % glycerol, 5 % beta-mercaptoethanol, 2 % SDS, 0.1 % bromophenol in ddH_2_O, pH 6.8) and stored at -80°C in aliquots.

### Spermidine detection via indirect ELISA

Human spermidine GENLISA^TM^ ELISA kit (Krishgen BioSystems, #KBH22460) was utilized for spermidine quantification. RAW264.7 macrophages were pre-treated with DFMO, DENSpm, spermine, and spermidine as mentioned above. Cells were collected into a fresh tube at 5×10^5^ cells/condition and pelleted by centrifugation for 5 min at 500×g. Cell culture medium was aspirated, and cells were resuspended in 200 µL PBS. Intracellular contents were released into PBS by three freeze-thaw cycles. Cellular debris was removed by centrifugation for 10 min at 800×g. Samples or standard dilutions were added in a 100 µL volume to the pre-coated wells with anti-Spd antibodies. The samples were incubated for 80 min at 37°C, then washed 4 times. Biotinylated anti-spermidine antibody solution was added to samples according to manufacturer protocol and incubated for 50 min at 37°C. After 4 washes, streptavidin-HRP conjugate solution was added to the samples and incubated for 50 min at 37°C. After another 4 washes, TMB substrate was incubated with samples for 10 min at 37°C and inactivated with stop-solution. TMB substrate conversions were measured at 450 nm with a microplate reader (Tecan Infinite^®^ M Plex). Spermidine concentration in experimental samples was calculated using the standard curve.

### Trypan Blue cell viability assay

Cell density was determined using a hemacytometer. 10 μL of a 0.4 % trypan blue solution (Thermo Fisher Scientific, #15250061), was added to 10 μL of cells suspension. Number of viable cells were determined according to the manufacturer’s recommendations.

### WST-1 cell metabolic assay

WST-1 (Roche, #5015944001) cell proliferation assay was used to assess cell viability after drug treatments. Briefly, cells were treated and/or supplemented as described for the assay. WST-1 reagent was added at a 1:10 dilution to cell culture medium in 24-well plates and incubated at 37°C, 5 % CO_2_, and 95 % RH. Conversion of WST-1 to formazan was evaluated every 15 min, by measuring the absorbance at 450 nm and for reference at 660 nm. Measurements were performed until control samples reached an optical density (OD) of 1 to 1.2 on a microplate reader (Tecan Infinite^®^ M Plex). Data was normalized according to the manufacturers recommendations and compared as indicated.

### Quantification of phosphatidyl serine (PS)

To evaluate the translocation of PS, the GFP Certified Apoptosis/Necrosis Detection Kit (Enzo, #ENZ-51002) was used. RAW264.7 cell pretreatment and infection were performed as indicated. On the day of visualization, cells were washed once with PBS and overlaid with 1 % (v/v) of apoptosis detection reagent (AnnexinV-EnzoGold) in Binding Buffer, provided with the kit. Samples were incubated for 15 min at room temperature (RT), protected from light. After 15 min of incubation, the staining solution was removed. Nuclei staining was performed using Hoechst dye at a final concentration of 10 ng/ml in PBS. After 10 min of incubation at RT, protected from light, the staining solution was removed, and cells were washed with PBS. Cells were covered with a few drops of Binding Buffer to prevent cells from drying out. Cells were observed under an epifluorescence microscope (Zeiss). Three representative fields of view per sample were quantified and normalized to the total number of cells per field of view.

### SDS-PAGE and Western blotting

Tricine-Sodium Dodecyl Sulfate (SDS)-Polyacrylamide Gel Electrophoresis (PAGE) was used (57). Briefly, frozen protein lysates were thawed on ice and denatured at 90°C for 10 min on a heat block. Lysed samples were centrifuged (5 min, 15,700×g and 4°C) before loading onto gels. Proteins of interest up to 70 kDa were separated in 10% Tricine-SDS-PAGE gels, while 12% gels were used for larger proteins. SDS-PAGE was conducted at 80 V for 10 min followed by 120 V for 50 min (10% gels), 120 V for 65 min (12% gels). Proteins were transferred onto a 0.2 µm nitrocellulose membrane (Omnilab) in transfer buffer (25 mM Tris, 193 mM glycine, 20 % methanol in ddH_2_O) at 100 V (60 min for 10% gels, or 90 min for 12 % gels). Membranes were blocked with 5 % fat-free milk (Carl Roth) in PBS + 0.05 % Tween 20 (PBS-T) overnight at 4°C. The following steps were conducted at RT. After blocking, membranes were incubated with the respective primary antibody (5 % fat-free milk in PBS-T) for 1 h, then washed three times with PBS-T for 5 min; ™ by for 1 h with the secondary antibody (diluted in 5 % fat-free milk in PBS-T). Membranes were then washed 3 times with PBS-T for 5 min and treated with Enhanced Chemiluminescence (ECL) substrate reagents (Pierce ECL Western Blotting-Substrate, Thermo Fisher Scientific or #32209, or alternatively Western Lightning Plus-ECL, Perkin Elmer, #NEL103001). Bands were visualized using the ImageQuant™ 800 western blot imaging system (Amersham).

### HNoV stool samples

HNoV isolates were obtained from stool samples from norovirus positive patients at the University Clinic of Schleswig-Holstein (UKSH) and collected in accordance with the study protocol approved by the Ethics committee of the University of Lübeck (approval number: 20-094). HNoV was genotyped by phylogenetic analysis of ORF1 and ORF2 sequences as previously described (58, 59). HNoV GEs were determined by RT-qPCR as previously described (60, 61). Short term stocks were prepared from stool filtrates (1 % solution in Opti-MEM, filtered through 0.22 µm membrane) and stored as aliquots at -80°C. HNoV inoculum was freshly prepared as indicated and directly used for infection of HIE.

### HIE maintenance and infection with HNoV

HIE line HT-124 (61) was maintained at 37°C, 5 % CO_2_, 95 % RH in basal membrane extract (BME, Matrigel Matrix, Corning, #354234) with growth medium (IntestiCult^™^, STEMCELL Technologies, #06010). HIEs were passaged every 7 days with a 1:2 split ratio and medium replacement every other day. Differentiation was by Wnt3a withdrawal, switching to standard differentiation medium (Advanced DMEM-F12, 1 % P/S, 1 % HEPES, 5 % Noggin conditioned medium, B-27 (Invitrogen, #17504-044), N-2 (Invitrogen, #17502-048), 50 ng/ml murine EGF (Thermo Fisher Scientific, #PMG8043), 1 mM N-acetyl cysteine (Sigma-Aldrich, #A9165), 10 nM Leu15-Gastrin (Sigma-Aldrich, #G9145), 500 nM A 83-01 (Tocris, #2939)) or putrescine-free differentiation medium (Advanced DMEM-F12, 1 % P/S, 1 % HEPES, 5 % Noggin conditioned medium, 50 ng/ml murine EGF (Thermo Fisher Scientific, #PMG8043), 1 mM N-acetyl cysteine (Sigma-Aldrich, #A9165), 10 nM Leu15-Gastrin (Sigma-Aldrich, #G9145), 500 nM A 83-01 (Tocris, #2939)) for 7 days with medium replacement every other day.

On the day of infection, 3D-HIEs were collected in washing medium (Advanced DMEM-F12 1 % P/S, 1 % HEPES) by scraping and pelleted at 90×g for 4 min at 4°C. HIEs were resuspended in differentiation medium and transferred to 1.5 ml tubes. HNoV filtrates were added at estimated MOI of 1 (based on GEs) to HIEs with 500 µM bile acid (glycochenodeoxycholic acid, GCDCA, Sigma-Aldrich, # G0759). Each sample contained an estimated 40 - 60 matured spheres in a final volume of 200 µL/tube. Virus and HIEs were incubated for 1 h at 37°C with occasional mixing. HIEs were washed three times with washing medium and pelleted (100×g for 2 min at 4°C). Spheres were plated in 30 µL/sample of BME into a pre-warmed 24-well plate. Plates were left for 10 min at 37°C to ensure polymerization of the BME. HIEs were supplemented with differentiation medium including 2 µM of Ruxolitinib (Sigma-Aldrich, # AMBH324A54EE) and 500 µM bile acid GCDCA. HIEs were harvested in 400 µL/well of TRIzol™ reagent (Zymo Research Europe GmbH), immediately (0 hpi), or 48 hpi.

### Titration of HNoV

HNoV titers were determined as previously described (51, 61). Briefly, total RNA was extracted from samples using Direct-zol RNA MiniPrep kit (Zymo Research Europe GmbH, # R2073). HNoV GEs were determined using the Luna Universal Probe One-Step RT-qPCR Kit (NEB) using specific primers and probe targeting the ORF-1/ORF-2 junction (forward: AGCCAATGTTCAGATGGATG, reverse: TCGACGCCATCTTCATTCAC, TaqMan probe: TGGGAGGGCGATCGCAATCTGGC). A synthetic dsDNA fragment (gBlock™, Integrated DNA Technologies IDT) corresponding to the amplified cDNA was used to generate a standard curve for quantification of HNoV genome eqs.

### Statistical analysis

Graphs were generated and statistically analyzed using GraphPad Prism, version 10 (GraphPad Software Inc., La Jolla, CA, 2023). For comparisons among multiple groups, a parametric ANOVA was applied when data met normality assumptions. The non-parametric Kruskal–Wallis test was used when normality could not be confirmed using the Shapiro–Wilk test.

## Results

### MNV is sensitive to PA depletion

PAs are required for a number of distinct viral families, including enteric viruses and other RNA viruses (37). To examine the role PAs in norovirus infection, we sought to determine MNVs sensitivity to PA depletion (**Figure 1**) using well established PA inhibitors (62).

Firstly, we employed N¹,N¹¹-diethylnorspermine (DENSpm), an activator of spermine/spermidine N¹-acetyltransferase 1 (SAT1). SAT1 catalyzes the acetylation of spermine and spermidine, promoting their degradation and excretion, thereby depleting them from the cell (**Figure 1a**). Three different cell lines were used, Huh7^mCD300lf^, constitutes a recombinant epithelial cell, expressing the murine mCD300lf receptor for susceptibility (51), BV-2 and RAW264.7 are microglia-like and macrophage-like monocyte-derived immune cell lines. All cells were pretreated with 100 µM DENSpm for 24 h prior to infection. Following pretreatment, cells were infected with MNV-1 at MOI 0.1 for 1 hour on ice, then shifted to 37 °C in the continued presence of DENSpm, with and without supplemented spermine or spermidine. Viral titers were then quantified by plaque assay 24 hpi. The data showed that DENSpm treatment significantly impaired MNV-1 infection across all tested cell lines (**Figure 1b-d**), with significant reductions of 2–3 logs in BV-2 and RAW264.7 cells, and up to 5 logs in Huh7^mCD300lf^ cells. Importantly, supplementation with 100 µM spermine or spermidine restored viral titers close to untreated controls (**Figure 1b-d**).

Secondly, we employed difluoromethylornithine (DFMO), a well-tolerated and specific inhibitor of ornithine decarboxylase 1 (ODC1) (**Figure 1a**). DFMO blocks the rate-limiting step in PA biosynthesis, the conversion of ornithine to putrescine, resulting in reduced levels of putrescine, spermidine, and spermine. Huh7^mCD300lf^, BV-2, and RAW264.7 cells were treated with 500 µM DFMO for 4 days prior to infection (**Figure 1e-g**). Cells were inoculated with MNV-1 at a multiplicity of infection (MOI) of 0.1 for 1 h on ice, followed by an incubation for 24 h at 37 °C in the continued presence of DFMO, with or without supplementation of spermine or spermidine. Similar to DENSpm, DFMO treatment led to significant inhibition of MNV infection across all tested cell lines by 24 hpi (**Figure 1e-g**), highlighting the role of PAs as critical host factors for MNV infection in both immune and epithelial cell types.

Similar data was also obtained for the MNV strain CR3 in Huh7^mCD300lf^ cells (**Figure A1**), showing that this is not restricted to MNV-1. Here PA-depletion treatment led to an almost 5-log reduction in infection 3 dpi. Unlike MNV-1, CR3 causes persistent infections in mice (17), providing a first link between MNV persistence and PAs.

To support the selected concentrations for both DENSpm and DFMO, dose–response experiments were performed in RAW264.7 cells, demonstrating a concentration-dependent inhibition of MNV infection (**Figure A2**).

Metabolic activity and cell viability were assessed using a WST-1 and a trypan blue exclusion assay and confirmed no negative effect on cell health during PA depletion (**Figure A3**). Notably, the addition of spermine or spermidine to untreated (PA-intact) cells did not alter viral titers (**Figure A4**), suggesting that baseline PA levels in standard culture conditions were adequate to support infection. Interestingly, supplementation with spermine or spermidine not only rescued viral replication in PA-depleted cells but increased MNV-1 titers beyond baseline levels (**Figure 1e-f**), which may be linked to the increase in metabolic activity observed in the WST-1 assay (**Figure A3**). Although the specificity and working concentrations of DENSpm and DFMO is well established (37), we confirmed a decrease in intracellular spermidine upon treatment in RAW264.7 cells with DFMO and DENSpm, as well as the restoration of intracellular spermidine upon supplementation in the cell culture medium (**Figure A5**). While serum was postulated to contain PAs and PA-catabolizing enzyme (63), we detected no spermidine in our serum and observed normalization of spermidine levels in PA-restricted cells, indicating no negative impact on exogenous supplemented PAs in presence of serum. While it is possible that levels were below the detection limit of the ELISA, or our cell culture serum contains PAs other than spermidine that we are unable to detect with this assay, they are unlikely to significantly impact our assay conditions.

In conclusion, our data demonstrates that PA depletion significantly blocks MNV infection across multiple cell types in a reversible and strain-independent manner.

### MNV attachment to cells is not affected by PA depletion

Previous reports have shown that other enteric RNA viruses rely on PAs for efficient binding and entry into host cells (64, 65). PAs regulate the expression of the transcription factor SREBP2, which is required for intracellular cholesterol biosynthesis (44). Reduced SREBP2 activity leads to decreased cholesterol content in cellular membranes, which could affect viral entry. Since MNV entry is cholesterol-dependent (66, 67), we investigated, whether PA depletion affects MNV-1 attachment to host cells. Huh7^mCD300lf^, BV-2 cells, and RAW264.7 cells were cultured in the presence of 500 µM DFMO for 4 days to deplete PAs (**Figure 2**). Cells were then inoculated with MNV-1 at MOI 5 and incubated on ice for 1 hour to allow viral binding without internalization. The unbound virus was removed by washing, and cells were harvested in TRIzol for total RNA extraction. Viral genome equivalents were quantified by RT-qPCR. PA depletion did not impair MNV-1 attachment to any of the tested cell lines (**Figure 2**). Also, the addition of spermine had no effect on viral binding to host cells. This suggests that in case of MNV-1 PA-depletion or PA-supplementation does not affect MNV-1 attachment.

**Figure 2.**
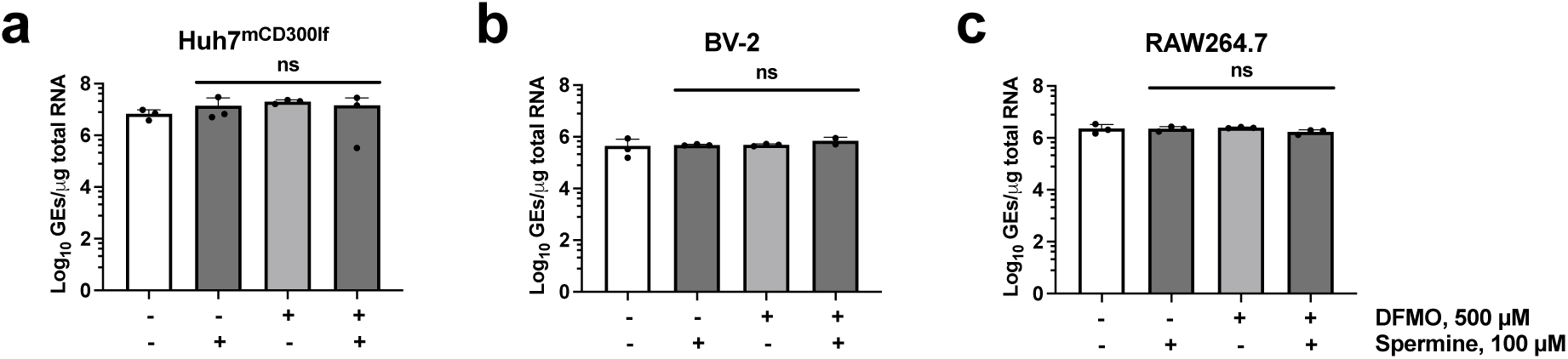
PA depletion does not alter MNV-1 attachment to cells. Huh7^mCD300lf^, BV-2, and RAW264.7 were pretreated with 500 μM DFMO for 4 days. Cells were inoculated with MNV-1 at an MOI of 5 on ice in the presence of DFMO and 100 µM spermine. After 1 h incubation to allow for virus attachment, cells were washed and harvested for total RNA extraction. MNV titers were quantified by RT-qPCR. Data represent means ± standard deviation (SD) of three independent biological assays. Individual data points are shown. Statistical significance was assessed using the Kruskal–Wallis test and shown compared to non-treated/infected: ns, not significant.

### PA depletion does not impact MNV-1 genome levels

For several flaviviruses, including Zika and Chikungunya virus, PAs are important for RNA genome replication and packaging and have been shown to promote the formation of membrane-associated replication complexes (68) (69). To investigate the effect of PA on MNV-1 genome replication, we pretreated RAW264.7 cells for 4 days with DFMO and infected cells with MNV-1 at an MOI of 5. Lysates were then harvested 16 hpi, with genome equivalents being quantified by RT-qPCR (**Figure 3a**) and infectious units by plaque assay (**Figure 3b**) from the same sample.

**Figure 3.**
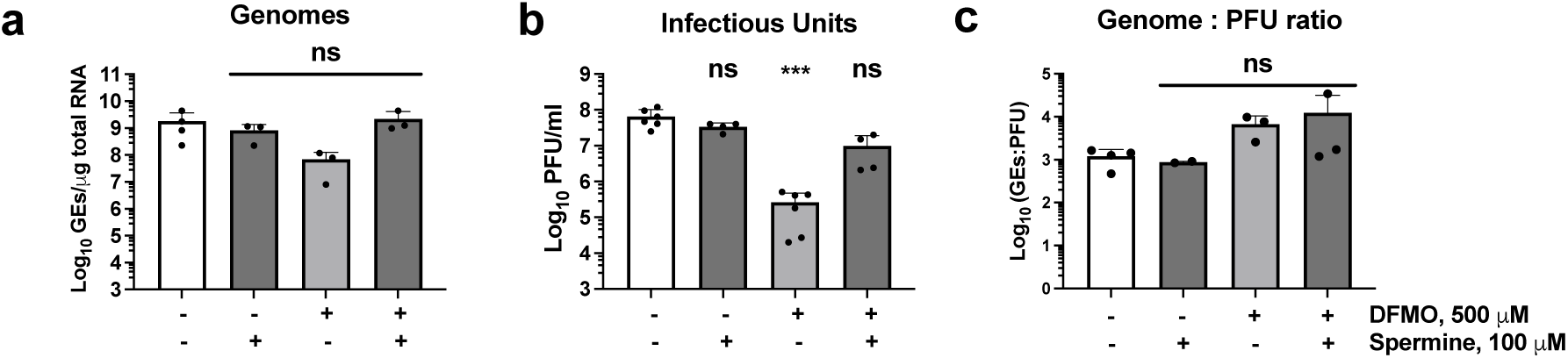
Impact of PA depletion on genome equivalents versus infectious units in RAW264.7 cells. RAW264.7 cells were treated with 500 μM DFMO for 4 days to deplete PAs, then infected with MNV-1 at an MOI of 5. Inoculated cells were supplemented with DFMO and/or 100 µM Spm. Cells were harvested by freezing at 16 hpi for quantification of viral genome via RT-qPCR **(a)** and infectious particles via plaque assay **(b)**. The genome to PFU ratio is depicted **(c)**. Data represent means ± standard deviation (SD) of at least three independent biological assays with two technical repeats for the plaque assay. Individual data points are shown. Statistical significance was assessed using the Kruskal–Wallis test and shown compared to non-treated/infected: ns, not significant; ***p≤0.001.

No significant effect was observed for MNV genome replication by PA depletion, while significant reduction was observed infectious units. The genome to PFU ratio in PA-depleted cells was slightly increased, suggesting that decline in genomes is insufficient to account for the decline in viral titers (**Figure 3c**). This prompted us to investigate whether PA impacts other steps in the infectious cycle after genome replication to contribute to the overall decline in viral titers.

### PA depletion prevents MNV-1 release

To investigate a potential release phenotype, infectious units in cell and supernatant fraction were determined independently between 8 to 48 hpi (**Figure 4**).

**Figure 4.**
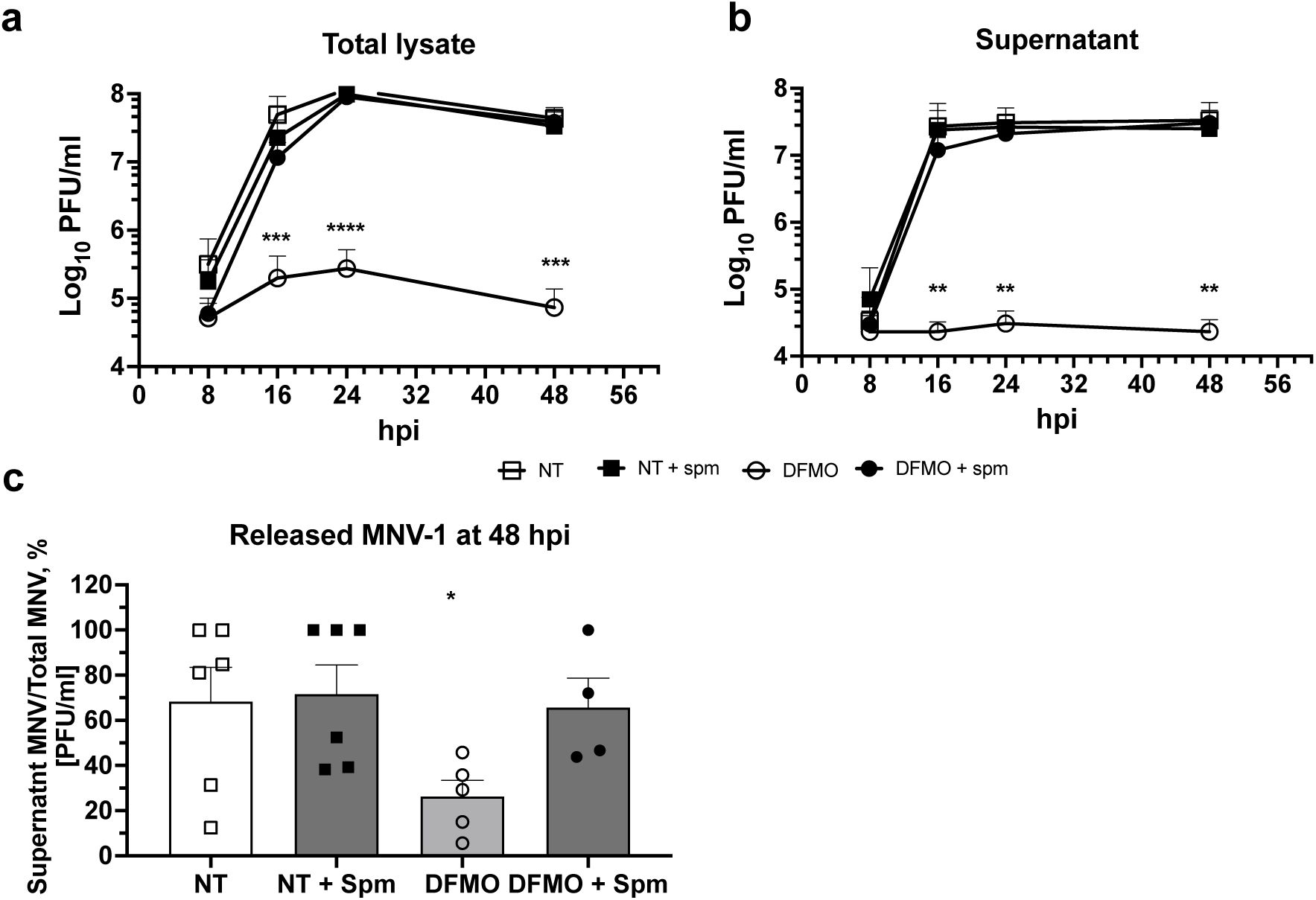
PA depletion impairs MNV-1 release. RAW264.7 cells were pretreated with 500 μM DFMO for 4 days to deplete PAs, then infected with MNV-1 at an MOI of 5. Infected cells were supplemented with/without DFMO and/or 100 μM spermine (Spm). Supernatant was collected separately, and virus titers were determined by plaque assay. Data represent means ± standard deviation (SD) of three independent biological replicates and two technical repeats for each condition. Statistical significance was assessed using a non-parametric Kruskal– Wallis test and shown compared to non-treated (NT): *p≤0.05; **p≤0.01; ***p≤0.001.

RAW264.7 cells were pretreated with 500 μM DFMO for 4 days and infected with MNV-1 on ice at MOI of 5. Following infection, the inoculum was removed, and cells were supplemented with medium containing DFMO, and/or 100 µM spermine and incubated for 8 to 48 h. To determine if virus release was affected, lysate and supernatants were collected separately, and infectious units were determined respectively for each time point. Infectious units were significantly and consistently diminished upon DFMO treatment in both lysate and supernatant fractions, showing that PA depletion impacts MNV-1 infection early during infection. Supplementation with spermine fully restored infection at all time points. In DFMO-treated lysates, titers increased between 8 and 24 hpi, which was not observed in the corresponding supernatants. Collectively, these findings demonstrate that PA depletion inhibits release of virions from infected cells.

### PA restriction prevents MNV-induced cell death

Another prominent phenotype observed was an almost complete loss of the cytopathic effect (CPE) in infected but PA-depleted cells (**Figure 5a**). Even at high MOI infection 48 hpi, DFMO-treated cells showed no CPE (**Figure 5a** panel v) compared to infected but non-treated cells (panel iv). This phenotype was again completely reverted by adding spermidine (panel vi) compared to infected and DFMO-treated (panel v).

**Figure 5.**
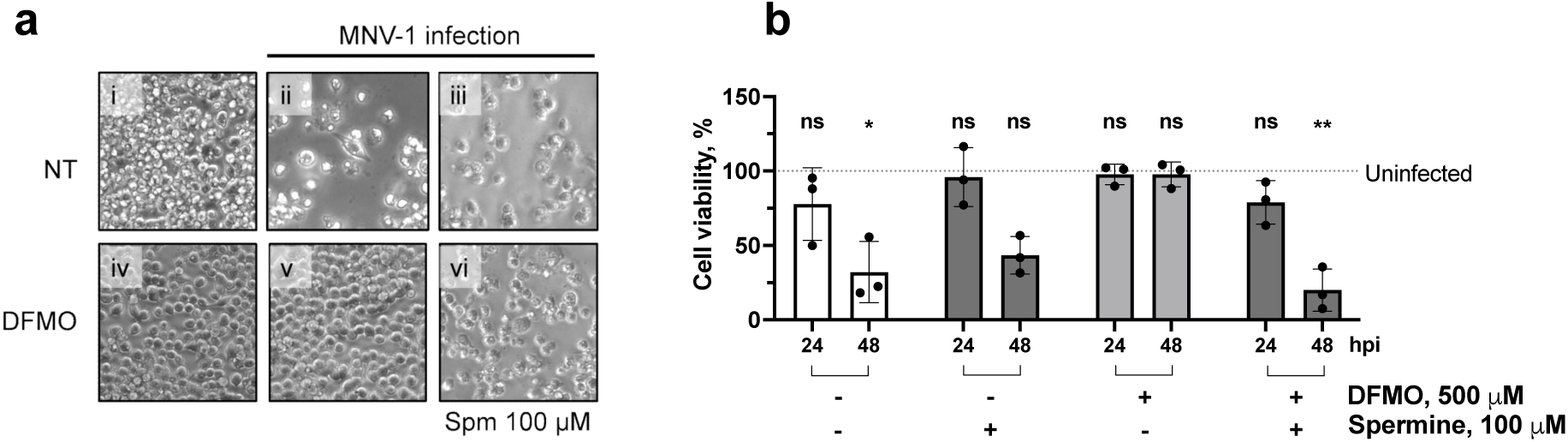
Loss of virus-induced cytopathic effect in PA depleted murine macrophages. RAW264.7 cells were pretreated with 500 μM DFMO for 4 days, and subsequently infected with MNV-1 at MOI of 5 on ice for 1h. After infection, the inoculum was removed, and cells were reconstituted with medium containing DFMO and/or 100 μM spermidine. MNV-induced CPE was analyzed 48 hpi by bright-field microscopy **(a)** and cell viability was determined by WST-1 assay at 24 and 48 hpi and shown as percent viable compared to the respective uninfected control **(b)**. Data are presented as means ± standard deviation (SD) from three independent biological experiments. Statistical significance was assessed using a non-parametric Kruskal–Wallis test and shown compared to the corresponding non-treated (NT)/non-infected sample: Non-significant statistical comparisons (p>0.05) are not shown for clarity; *p≤0.05.

To quantify this effect on cell viability a WST-1 assay was performed showing that about 30% of the infected cells died 24 hpi and up to 70 % by 48 hpi (**Figure 5b**). However, no change in cell death was observed in infected and DFMO-treated cells. Here, the cell death was again completely restored when spermine was supplemented. These data suggest that PAs are necessary for MNV-1 induced cell death.

### PA depletion prevents MNV-induced PARP cleavage linked to PI3K signaling and apoptosis

Recently it was shown, that MNV infection promotes the cleavage of apoptotic caspase-3 and PARP and that apoptosis is important for efficient MNV replication and viral release (27). Because a protective effect of PA on apoptosis has been widely known (45, 70), we hypothesized that the lack of CPE in PA-depleted cells (**Figure 5**) may be linked to the inability of the virus to induce apoptosis. Because, we struggled to consistently detect caspase-3 cleavage in MNV-1 infected cells, we investigated PARP cleavage, downstream of caspase-3 cleavage, as a marker for virus-induced apoptosis (**Figure 6**). Analyses were performed in RAW264.7 and Huh7^mCD300lf^ cells that were infected with MNV-1. Cells were depleted for PAs using DFMO and reconstituted at first using spermine supplementation. Similarly to the reported observations in immortalized bone marrow-derived macrophages (iBMDM) (27), our data confirmed that MNV-1 infection led to a loss of uncleaved PARP in PA-competent RAW264.7 and Huh7^mCD300lf^ cells. Interestingly, in DFMO-treated and infected cells, no virus-induced loss of PARP occurred 24 hpi (**Figure 6**), suggesting that in PA-depleted cells PARP is not cleaved and apoptosis inhibited.

**Figure 6.**
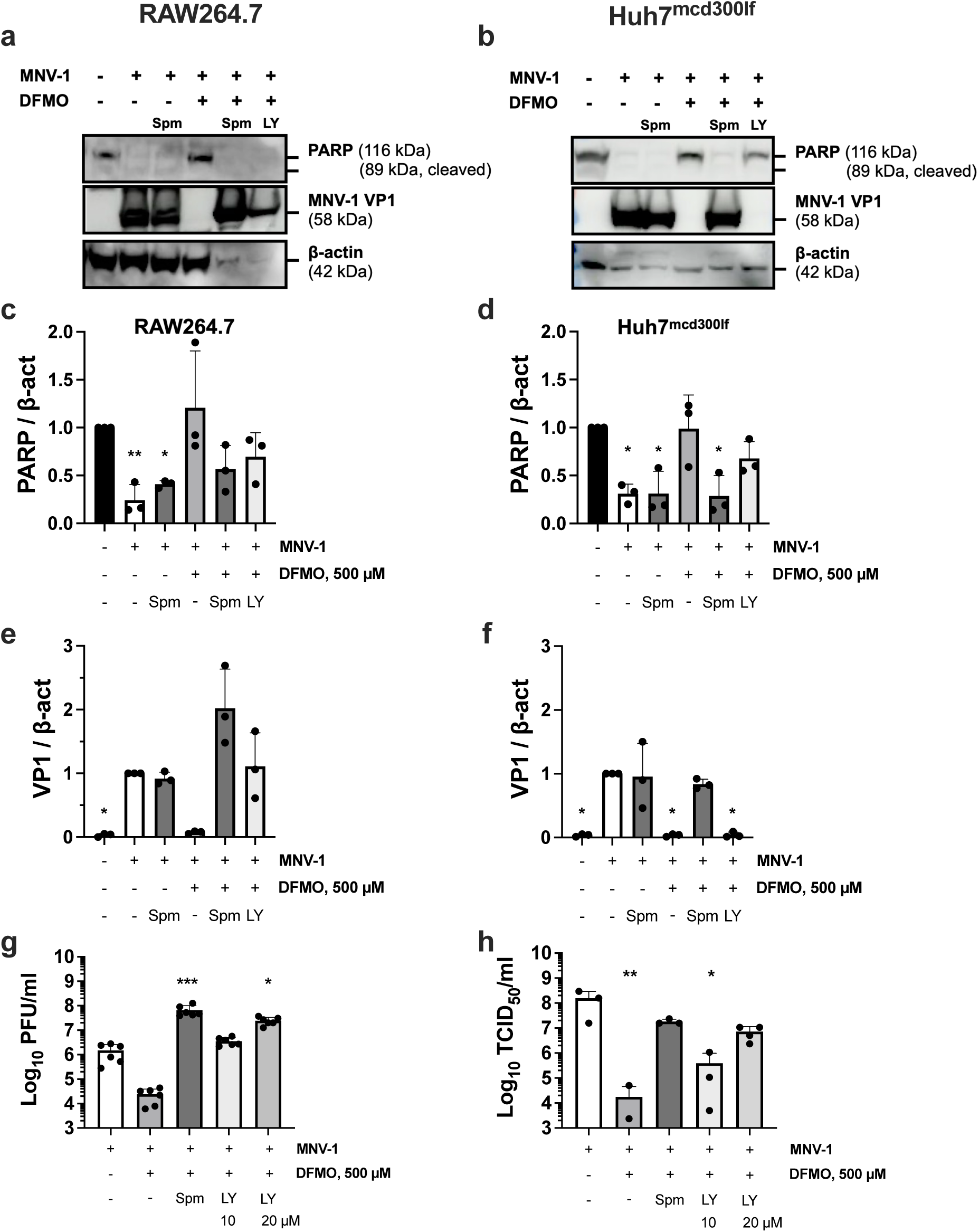
PA depletion prevents MNV-1 induced PARP cleavage and infection, rescuable by PA reconstitution and PI3K inhibition. RAW264.7 and Huh7^mcd300lf^ cells were pretreated with 500 μM DFMO treatment for 4 days. Cells were then infected with MNV-1 at MOI 1 on ice for 1 h. Following infection, the inoculum was removed, and the medium was supplemented with/without DFMO, 10 μM LY294002 (LY) or 100 μM spermine (Spm). At 24 hpi, cell fractions were harvested for protein extraction and subsequently analyzed by western blotting to determine expression levels of MNV capsid protein VP1, PARP and β-actin (β-act). A representative western blot is shown for each cell line **(a, b)**. A quantification of PARP **(c, d)** and VP1 **(e, f)** was performed using ImageJ gel analysis and normalized to β-act. Data represent means ± standard deviation (SD) of three independent biological assays **(a-f)**. Individual data points are shown. For statistical analysis PARP expression was compared to the untreated and uninfected control, while VP1 was compared to untreated and infected samples. Virus titers were further determined by plaque assay **(g, h)**. Data are presented as means ± standard deviation (SD) from three independent biological and up to two technical repeats **(g-h)**. Individual data points are shown. Statistical significance was assessed in comparison to non-treated/infected. Statistical significance was assessed using the Kruskal–Wallis test: ns, not significant (omitted for clarity); *p≤0.05; **p≤0.01; ***p≤0.001.

Since inhibition of PI3K using LY294002 (LY) restored apoptosis induction by TNF-⍺ in DFMO treated intestinal epithelial cells (46), we next supplemented PA depleted cells with the PI3K inhibitor LY to restore MNV-1 induced apoptosis and PARP cleavage. Indeed, in infected PA-depleted RAW264.7 cells, LY restored PARP cleavage and MNV-1 infection similarly to spermine (**Figure 6**). Under these conditions, a loss of β-actin was observed, potentially reflecting the increased cell death. No PARP cleavage occurred in uninfected LY and spermine treated cells, showing that PARP cleavage is a consequence of the infection and not of PI3K inhibition or PA reconstitution (**Figure A6**). VP1 levels were also restored in the presence of spermine and LY, mirroring the rescued infection (**Figure 6g**). In Huh7^mCD300lf^ cells PI3K inhibition also showed a reduced PARP signal and partially restored MNV-1 infection. At an increased LY concentration (20 µM), infection was restored similarly to spermine (**Figure 6h**). Since we observed a complete block of viral release during the course of infection, we determined viral release at 48 hpi in the presence of both, spermine and LY. Here, both spermine and LY fully restored viral release in RAW264.7 cells (**Figure 7**).

**Figure 7.**
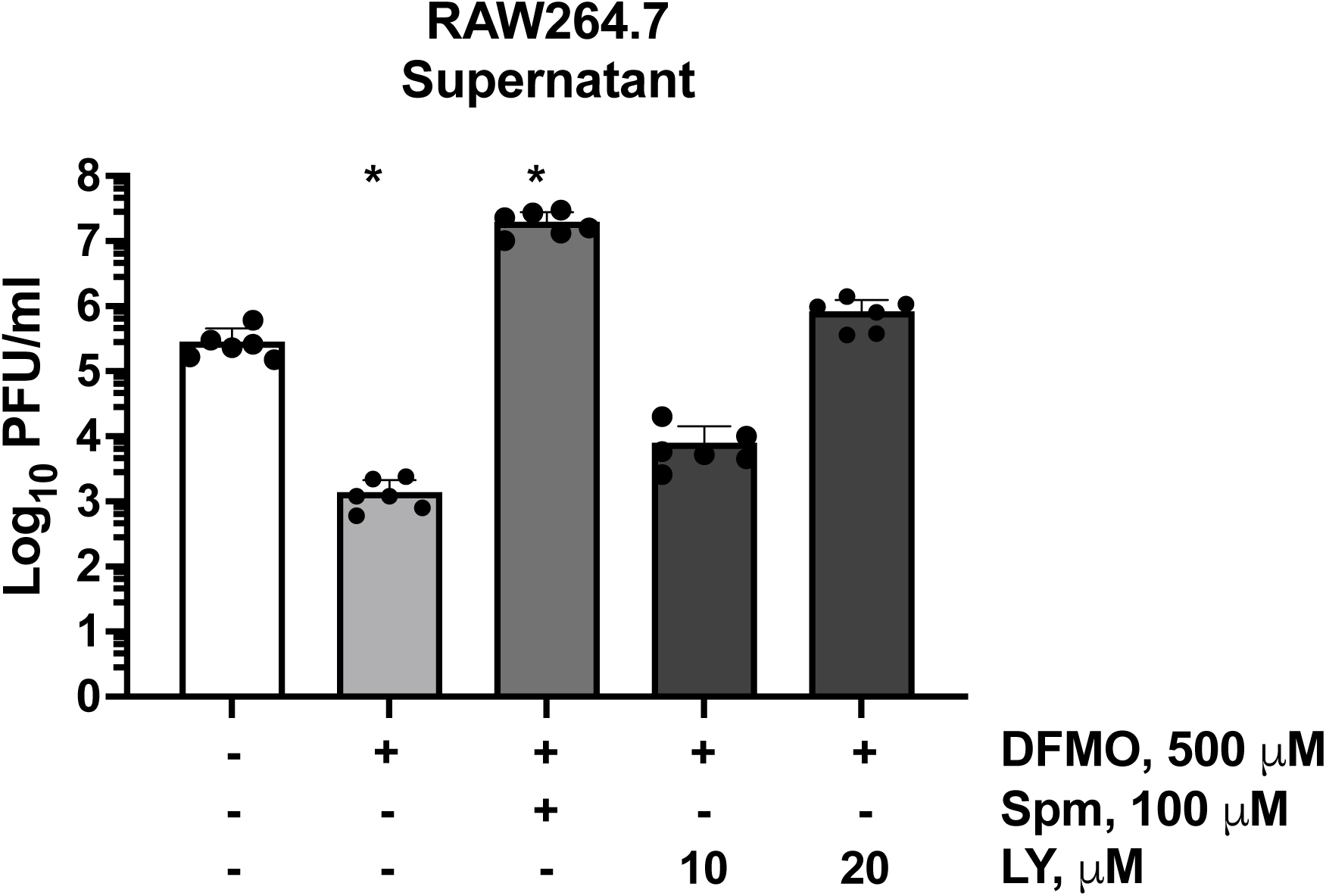
PI3K inhibition rescues MNV-1 release in PA-depleted cells. RAW264.7 cells were pretreated with 500 μM DFMO for 4 days to deplete PAs and infected with MNV-1 at an MOI of 0.1. Infected cells were supplemented with/without DFMO and/or 100 μM spermine (Spm), or 10 -20 µM LY294002 (LY). Supernatant was harvested 48 hpi, and virus titers were determined by plaque assay. Data are presented as means ± standard deviation (SD) from three independent biological repeats with two technical repeats. Individual data points are shown. Statistical significance was assessed using the Kruskal–Wallis test and shown as compared to non-treated: *p≤0.05.

In summary, loss of uncleaved PARP, indicative for apoptosis, correlated with successful infection and viral release and was dependent on PAs. PA depletion prevented MNV-1 induced PARP cleavage and promoted cell survival in a PI3K dependent manner.

### PA depletion inhibits MNV-induced apoptosis in murine macrophages

To address the PA depletion effect on MNV-induced apoptosis directly, we characterized the surface presentation of PS via Annexin V staining in MNV-1 infected macrophages (**Figure 8**).

**Figure 8.**
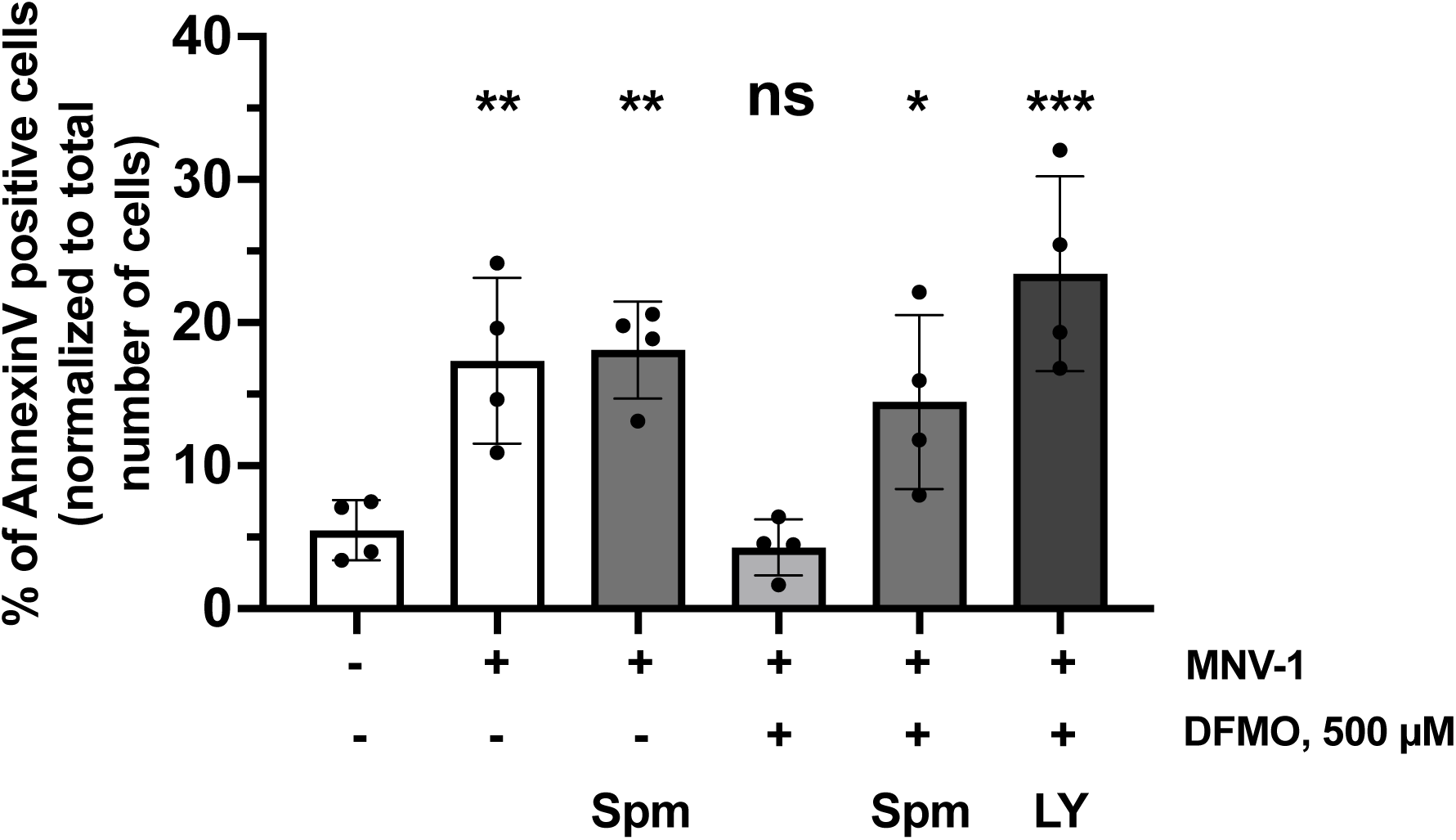
PS externalization in MNV-infected macrophages depends on PA availability. RAW264.7 cells were treated with 500 μM DFMO treatment for 4 days, then cells were infected at an MOI of 0.1 for 1h on ice. The inoculum was removed and replaced with a medium containing DFMO, 20 μM LY or 100 μM Spm. At 24 hpi, live cells were stained with AnnexinV to detect PS externalization and Hoechst for nuclei. PS-positive cells were quantified and normalized to total number of cells per field of view. Data represent means ± standard deviation (SD) of at least two independent biological assays. Individual data points are shown. Statistical significance was assessed in comparison to uninfected and untreated using ANOVA: ns, not significant; *p≤0.05; **p≤0.01; ***p≤0.001.

RAW264.7 cells were treated with DFMO and subsequently infected at an MOI of 0.1. During the infection, cells were supplemented as indicated and stained with Annexin V-AF at 24 hpi. MNV-1 infection induced phosphatidylserine (PS) externalization in approximately 17% of cells. In contrast, PA-depleted, MNV-infected macrophages showed PS externalization levels comparable to uninfected cells (∼5% Annexin V– positive). Supplementation with spermine or LY restored PS externalization in PA-depleted, infected cells to 14% and 20%, respectively, in line with our observations for loss of uncleaved PARP, infection, and viral release (**Figure 7**).

Collectively, these data show that MNV-1 relies on PA and PI3K signaling for apoptosis induction and successful infection and viral release.

### PA restriction ablates HNoV GII.4 infection in intestinal enteroids

To assess whether PAs play a broad role in norovirus infection, we investigated the impact of PA depletion using two distinct HNoV GII.4 stool-isolates (**Figure 9**). To assess the impact of PA restriction on HNoV infection, DENSpm was used to degrade PAs in HIEs, which resulted in full block of HNoV replication with partial titer rescue after supplementation of 100 µM spermine for both stool isolates (**Figure 9a-b**). Similar results were obtained with two distinct stools isolates, suggesting that the effect is not stool dependent. Since the HIE differentiation medium contains 150 µM putrescine, the use of DFMO, was not feasible, as putrescine supports downstream PA synthesis. We therefore used differentiation medium omitting the putrescine containing additives B27 and N2 (putrescine-free medium (B27-/N2-)) (**Figure 9c**). After 7 days of differentiation, HIEs were infected with GII.4 Den Haag #1 at MOI 1. After inoculation, putrescine-free medium was reconstituted with a mix 100 µM spermine and 100 µM spermidine. HNoV titers were quantified 48 hpi from whole cell samples via RT-qPCR. Again, depletion of putrescine from differentiation medium resulted in full restriction of HNoV infection and partial rescue supplementing PAs. Cell viability of HIEs were not significantly affected in either treatment (**Figure 9d**). However, differentiation in putrescine-depleted medium led to ≤30% loss in viability, which was partially restored by supplementation of PA mix. Consistent with our data on murine cells, an increase in metabolic activity was also observed only in DENSpm treated spermine supplemented HIEs.

**Figure 9.**
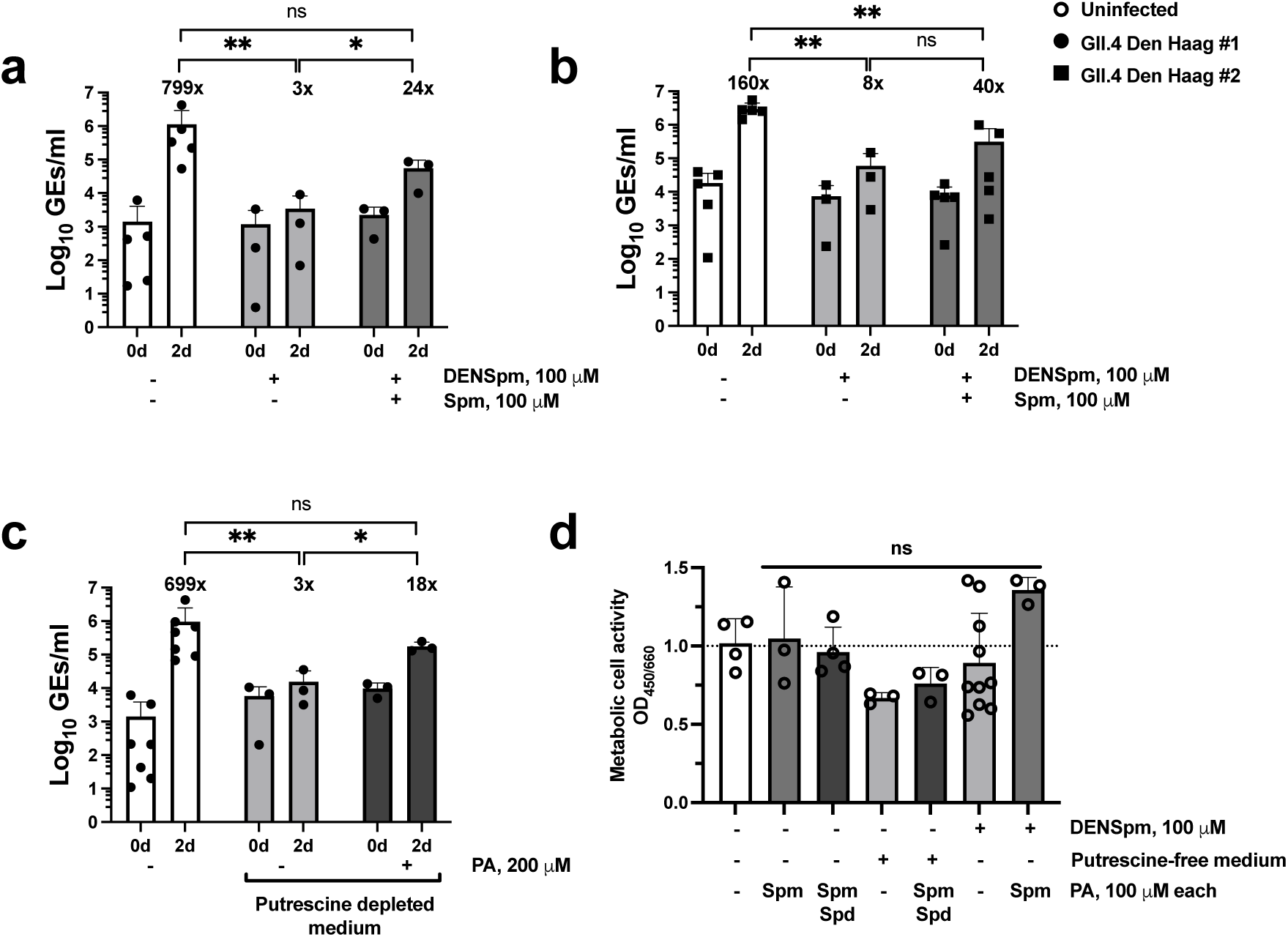
PA degradation inhibits HNoV GII.4 infection in HIEs. HIEs were differentiated in standard differentiation medium with 100 µM DENSpm additionally added 2 days before infection **(a-b)** or 7 days in putrescine-free medium (B27-/N2-) **(c)**. HIEs were infected with HNoV GII.4 stool isolates Den Haag #1 **(a, c)** or Den Haag, #2 **(b)**, at an approx. MOI of 1 for 1 h at 37°C. The inoculum was removed and replaced with standard medium with or without DENSpm and 100 µM Spm **(a-b)** or putrescine-free (B27-/N2-) medium with or without 200 µM PA mix (PA; 100 µM Spm + 100 µM Spd) **(c)**. HNoV GEs were determined by RT-qPCR at 48 hpi. Numbers above the bars indicate the fold-change in viral GEs at 2 days compared to 0 days. Viability of uninfected, treated HIEs were assessed using the WST-1 metabolic assay **(d)**. HIEs were treated as above, omitting only the viral inoculation. Data are presented as means ± standard deviation (SD) from a minimum of three independent biological assays. Individual data points are shown **(a-d)**. Statistical significance was assessed using ANOVA in comparison to untreated/infected **(a-c)** or just untreated **(d)**: ns, not significant; *p≤0.05; **p≤0.01.

Collectively, our data show that PAs also contribute to HNoV infection in HIEs.

## Discussion

Viruses have evolved to manipulate cellular signaling pathways to obtain essential biomolecules and to sustain optimal conditions for infection. PAs are ubiquitous cellular molecules that are present at a high micromolar concentration and play key roles in many cellular processes (39–43). They are important during infection by RNA and DNA viruses (36). For example, several RNA viruses, including other enteric viruses such as poliovirus, *CVB3,* and Enterovirus A71, rely on host-produced PAs for their replication (37, 62). Our study adds noroviruses to this group. We showed that restricting PAs in naturally susceptible murine immune cells and recombinant human epithelial cells significantly reduced MNV infection and that addition of PAs to the media could reverse this phenotype. Similarly, PA restriction in HIEs drastically inhibited HNoV GII.4 replication, suggesting that noroviruses exploit host-produced PAs to facilitate infection, similar to flaviviruses, coronaviruses, phleboviruses and enteroviruses (62).

Prior studies have linked PAs to virus attachment and replication of enteroviruses (71) and flaviviruses (62). Here, PAs were shown to affect the activity of transcription factor SREBP2, a crucial factor for cholesterol biosynthesis. Notably, cholesterol supplementation in PA-depleted cells rescued enterovirus CVB3 infection (44). Although MNV was shown to require cholesterol for cell entry (66), we found that PA-depletion did not impact MNV attachment to susceptible cells. However, the impact of PA supplementation on MNV entry and membrane cholesterol levels in PA-competent cells remains to be further addressed.

Many flaviviruses rely on host-derived PAs for viral genome replication and protein translation, we on the other hand observed no significant reduction in MNV genome replication in PA-depleted murine macrophages. In comparison, this was also the case for hepatitis B virus where it was shown that DFMO inhibits HBV replication by reducing HBc stability without affecting mRNA transcription or protein translation (72). In our case, PA instead facilitated MNV-induced apoptosis and promoted virion release.

Our findings indicate that noroviruses rely on host-derived polyamines to regulate apoptosis, as polyamine depletion markedly inhibited MNV-induced cell death. In PA-depleted macrophages, virus-induced apoptosis was reduced, evidenced by diminished PS externalization and PARP cleavage. Supplementation with spermine restored both MNV titers and apoptotic markers, highlighting the critical role of polyamines in MNV-mediated cell death. interestingly, PA-depleted murine macrophages and human recombinant epithelial cells remained susceptible to staurosporine-induced apoptosis, indicating that virus-associated apoptosis mechanisms were selectively affected by PAs restriction. Alongside, apoptosis activator LY restored virus-induced cell death and MNV titers. These observations suggest that PI3-kinase and downstream target proteins are involved in regulation of norovirus-induced cell death and apoptosis. LY also restored PS externalization and PARP cleavage during MNV infection, underscoring the importance of MNV-induced apoptosis in viral release. Together, these findings indicate that host PAs have a multifactorial role in MNV infection of murine macrophages, promoting virus-induced apoptosis and aiding virus release from infected cells.

While the PI3K/Akt molecular pathway is involved in many cellular processes (73), it is well established that activation of Akt protein via phosphorylation prevents apoptosis and prolongates cell survival (74, 75). MNV induces Akt phosphorylation at early time-points of infection to support cell survival during virus replication. Inhibiting Akt phosphorylation has been shown to interfere with optimum assembly of MNV virions and to promote early virus release, in a strain-dependent manner (14). Thus, precise Akt protein regulation appears critical for MNV egress. Restricting PAs is reported to lead to sustained Akt phosphorylation (46, 76, 77), which might alter infection dynamics and reduce apoptosis by inhibiting key events like PARP cleavage. Assessing the abundance of total Akt and phosphorylated Akt protein together with cleaved PARP protein would further clarify the role of Akt in norovirus-induced apoptosis.

Our study also has some limitations but already provides new avenues of research. While we assessed PAs restriction on MNV infection *in vitro* in several different cell types, extending these studies to primary murine macrophages and dendritic cells would further validate the role of PAs in MNV infection. The efficacy of PAs restriction against HNoV infection in the HIE model, however, already supports the biological significance of these findings in a non-transformed model. While we were able to exclude common PA targets such as viral attachment and genome replication and narrowed down the critical step between genome replication and viral release, the precise mechanism requires further investigation with particular emphasis on protein translation and virus assembly. Hypusination, where spermidine becomes covalently attached to eIF5A, has been linked to the ability of the host to translate proteins from certain mRNAs, particularly those containing polyproline stretches or other ribosome-stalling motifs that are otherwise difficult to translate efficiently. While there are no obvious polyproline stretches in the norovirus genome, hypusination could affect the ribosomal frameshifting that leads to the translation of ORF-3 (78).

Another open question is the impact *in vivo*. PA depletion significantly attenuated CVB3 and CHIKV replication in multiple organs of infected mice (62). PA supplementation in mice also enhanced herpes simplex virus 1 (HSV-1) infection (79). Mice treated with DENSP showed markedly reduced HSV-1 infections in brain and spleen. PAs also facilitate Alphavirus Infection *in vivo*, as shown for Sindbis virus (SINV) infection in the *Drosophila melanogaster* and zebra fish (*Danio rerio*) model, were animals were fed DFMO for 7 days prior to infection showed significantly reduced infections (68). These data show that PA modulation *in vivo* by supplementation or pharmacological depletion is an effective strategy against a broad range of PA dependent viruses.

Pancreatic ductal cells persistently infected with CVB3 lacked a robust virus-induced CPE and showed reduced PA levels (65). A similar situation might be possible for persistent MNV strains, such as CR3, which reside in an epithelial reservoir, by down-regulating IFN-λ signaling (20, 80). A direct feedback in which IFNs induce SAT1 expression, leading to PA depletion in a manner similar to DENSp, has also been described for Zika and CHIKV, leading to restricted infection (68). Whether PA depletion occurs in MNV-infected tuft cells or contributes to MNV persistence is an intriguing question that warrants further study.

In addition to intrinsic PA, gut bacteria are also a major source of PA in the intestine (41, 81, 82). The absence of commensal bacteria i.e. by antibiotic treatment would directly affect PA levels in the gut. Antibiotic treatment has been linked with reduced MNV infection in the gut and protection from disease (80, 83, 84), adding an extra layer of complexity that need to be taking into account for *in vivo* analysis and therapeutic approaches.

Understanding the role of host PAs in norovirus infection could guide antiviral therapy development. The PAs synthesis inhibitor DFMO is currently in clinical use for trypanosomosis (85) and demonstrates good tolerability (86). The PA analog DENSpm also has a good safety profile (87–89), although it still lacks clinical approval. Significant PA depletion requires 24 to 96 hours *in vitro*. While this indicates that PA inhibition may be less effective against acute norovirus infection, PA restriction may offer a promising therapeutic approach for patients with chronic norovirus infection, who suffer from prolonged diarrhea and are unable to clear infection. However, any therapeutic application would require considerations regarding external sources of PA in the gut, taking dietary PA uptake (90) and PA production by gut commensals (82) into account.

## Summary

In summary, our research highlights the role of PAs in norovirus infection. We demonstrate that restricting PAs reversibly inhibits MNV and HNoV infection by limiting virus release from various cell types. For MNV, we were able to show that PA depletion disrupts virus induced CPE linked to PARP cleavage and apoptosis. To the best of our knowledge, this is the first report associating PAs with virus-induced apoptosis. Our findings identify a novel target to exploit for intervention against norovirus infection open to already licensed pharmacological approaches such as DFMO, as well as dietary and targeted microbiome modulation approaches to deplete PAs in the gut.

## Acknowledgements

CM was supported by the Marie-Skłodowska Curie Actions Global Fellowship (GA-841247). MC was supported by the emergency aid programme to support refugee academics from Ukraine at the University of Lübeck.

## Notes

### Competing Interest Statement

The authors have declared no competing interest.

