## Supplemental Figures for "Polyamine depletion inhibits norovirus infection by blocking virus-induced apoptosis"

### Appendix

***Figure A1***

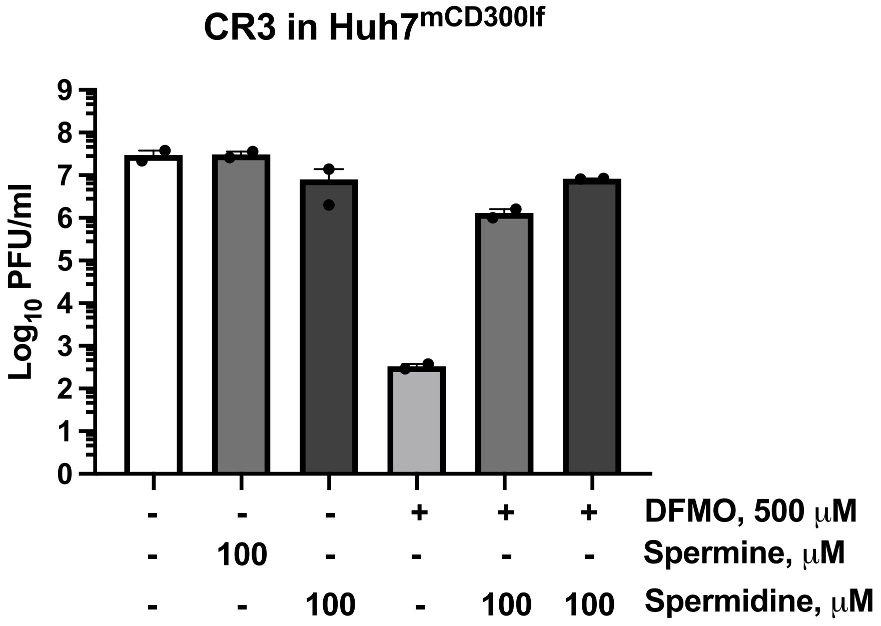


***Figure A1. PA depletion inhibits MNV CR3 infection in epithelial cells.*** *Huh7[mCD300lf] were pretreated with 500 μM DFMO for 96 h. Cells were inoculated with MNV CR3 at MOI 0.1 for 1 h at 4°C. The inoculum was removed and replaced with culture medium (-) or supplemented with DFMO and/or 100 μM of spermine or spermidine. Virus titers were determined 24 hpi by plaque assay. Data represent means ± range of two independent biological assays. Individual data points are shown.*

*Figure A2*

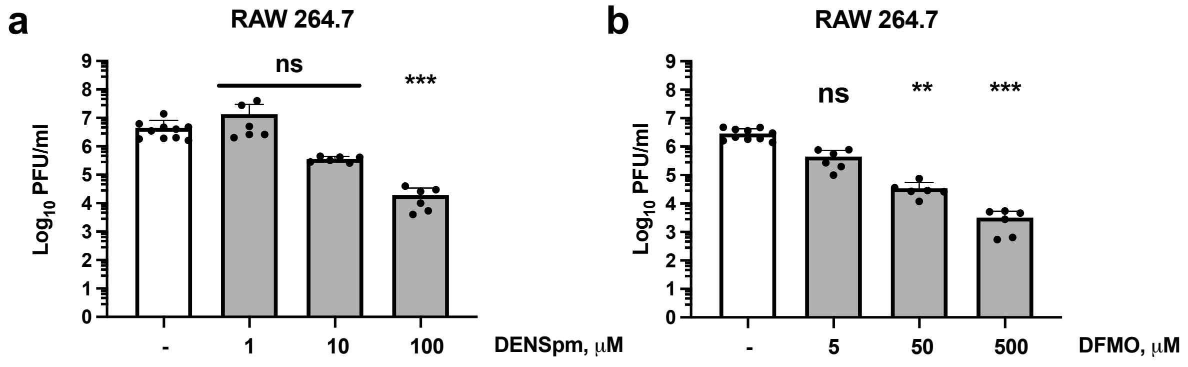


***Figure A2. Dose-dependent inhibition of MNV-1 infection by DENSpm and DFMO.*** *RAW264.7cells were pretreated with increasing concentration of DENSpm for 18 h or DFMO for 96 h. Cells were inoculated with MNV-1 at MOI 0.1 and for 1 h at 4°C. The inoculum was removed and replaced with culture medium (-) or medium supplemented with DENSpm or DFMO. Virus titers were determined 24 hpi by plaque assay. Data represent means ± standard deviation (SD) of three independent biological assays with two technical repeats each. Individual data points are shown. Statistical significance was assessed using the Kruskal–Wallis test compared to non-treated/infected: ns, not significant; **p≤0.01; ***p≤0.001.*

#### *Figure A3*

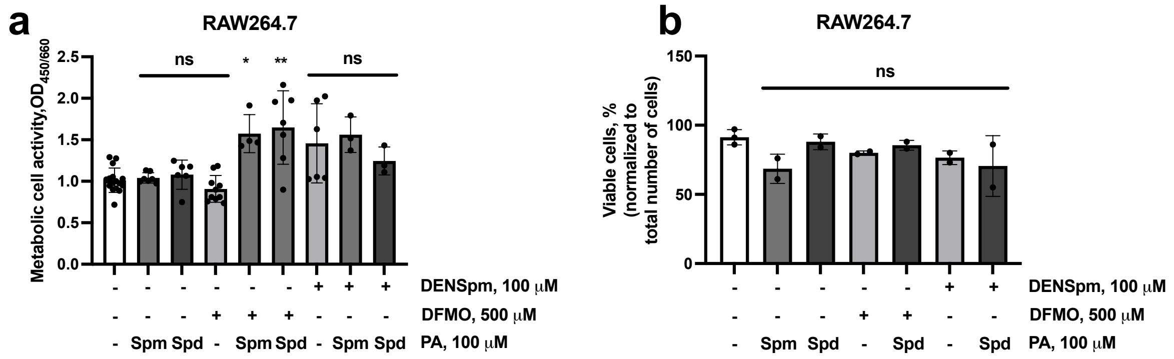


***Figure A3. PAs and PA-modulating drugs do not affect RAW264.7 macrophage viability or metabolic activity.*** *RAW264.7 cells were pretreated with 100 µM DENSpm for 24 hours or 500 µM DFMO for 96 hours. Following pretreatment, the medium was replaced with either control (mock) or supplemented with DENSpm, DFMO, 100 µM Spd, or 100 µM Spm. After 24 hours of Spm or Spd supplementation, metabolic activity was assessed using a WST-1 assay (a), and cell viability (b) was determined by trypan blue exclusion. Data represent means ± standard deviation (SD) of three independent biological assays. Individual data points are shown. Statistical significance was assessed using the Kruskal–Wallis test compared to non-treated: ns, not significant; *p≤0.05; **p≤0.01.*

*Figure A4*

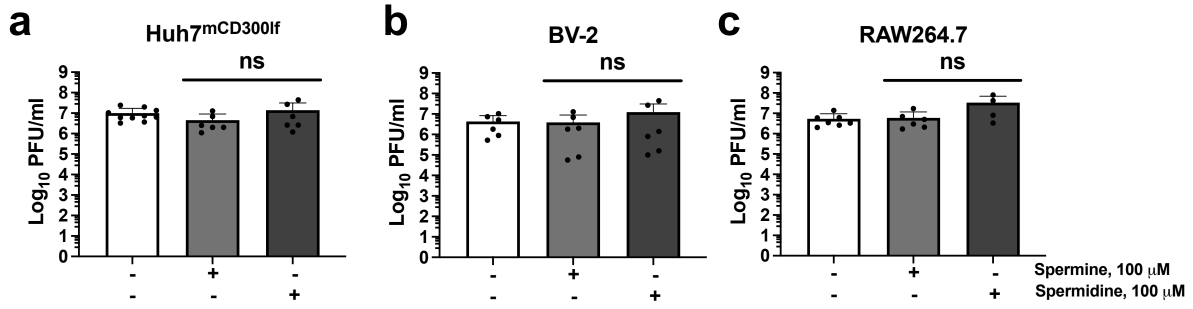


***Figure A4. PA supplementation is not affecting MNV-1 infection in immune and epithelial cells.*** *Huh7$^{mCD300lf}$(A), BV-2(B) and RAW264.7(C) cells were infected with MNV-1 at a MOI of 0.1 for 1 h on ice. The inoculum was removed and medium containing 100 µM Spm or 100 µM Spd were supplemented to cells. At 24 hpi, samples were harvested for titer quantification via plaque assay. Data represent means ± standard deviation (SD) of three independent biological assays with two technical repeats. Individual data points are shown. Statistical significance was assessed using the Kruskal–Wallis test compared to non-treated: ns, not significant.*

#### *Figure A5*

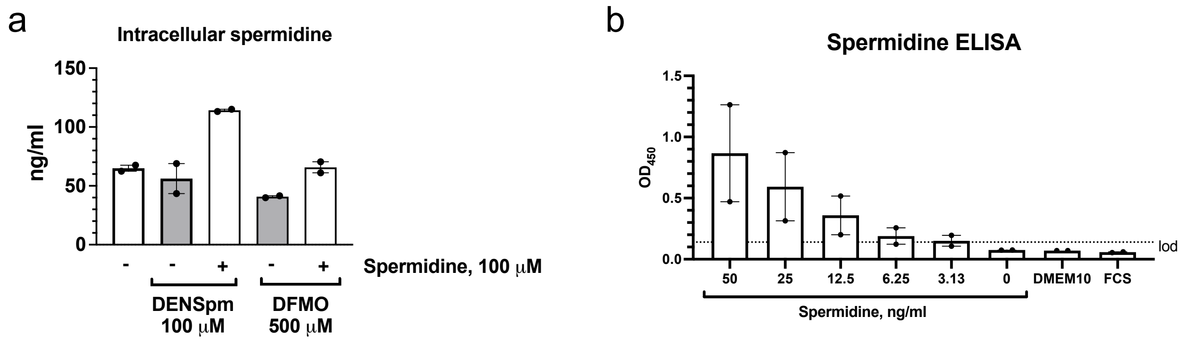


***Figure A5. PA degradation and depletion reduce intracellular spermidine concentration in RAW264.7 macrophages.*** *(a) RAW264.7 were pretreated with 100 μM DENSpm for 24 h or 500 μM DFMO for 96 h. Spermidine were supplemented to cells 6 h prior to harvesting samples by freezing. Efficacy of PA restriction drugs was assessed by measuring intracellular spermidine via indirect ELISA assay and normalizing to mock treated condition. (b) Spermidine content of cell medium components was also measured via indirect ELISA assay. Data represent means ± range of two independent biological assays. Individual data points are shown.*

*Figure A6*

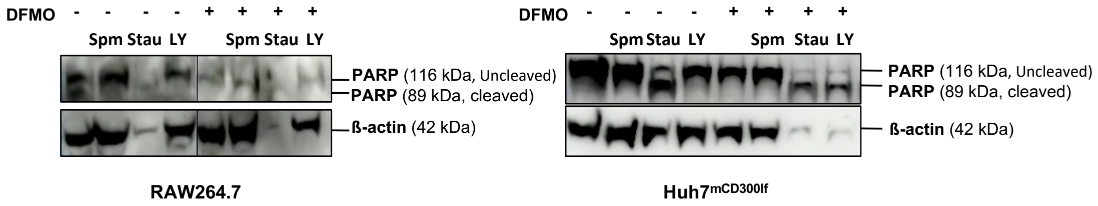


**Figure A6. Effect of PA and PA modulation drugs on apoptosis induction.** Uninfected murine macrophages RAW264.7 (left panel) and human recombinant epithelial Huh7$^{mCD300lf}$ cells (right panel) were pretreated with 500 µM DFMO for 4 days. Following treatment, the medium was replaced with a fresh medium containing DFMO, 20 µM LY, 100 µM Spm and 2 µM staurosporine. At 24 hours post treatment, cell fractions were harvested for protein extraction and subsequently analyzed by western blotting to determine expression levels of PARP. β-actin was used as a reference protein.
